# Ultra processed microbiome: Effects of dietary Stable Carbonyl adducts (SCars) on developing microbiota of mice

**DOI:** 10.64898/2026.09.17.752391

**Authors:** David Turner, Andrea Ottesen, Brandon Kocurek, Jackson Lane, Victoria J. Findlay

**Affiliations:** Department of Surgery, Virginia Commonwealth University, Richmond VA, USA; Massey Comprehensive Cancer Center, Virginia Commonwealth University, Richmond VA, USA; Center for Veterinary Medicine, Food and Drug Administration, Laurel, MD 20708, USA

**Keywords:** murine microbiome, stable carbonyl adducts (SCars), advanced glycation end-products (AGEs), metagenomics, ultra-processed food (UPF)

## Abstract

Consumption of ultra-processed food (UPF) is increasing and is causally linked with noncommunicable disease onset, however specific compounds within UPFs that influence disease risk remain poorly defined. The acronym ‘SCars’, for Stable Carbonyl Adducts, is introduced herein as an umbrella term encompassing advanced glycation end-products (AGEs), advanced lipoxidation end-products (ALEs), and other carbonyl-derived modifications. These molecules arise as products of spontaneous carbonyl chemistry, a reaction accelerated by industrial food processing. Dietary SCars have been associated with metabolic dysfunction; however, their impact on microbiome composition during windows of developmental vulnerability has not been well studied. Puberty in mice represents a period of developmental plasticity during which the gut microbiome is susceptible to dietary exposures. We tested whether transient exposure to high levels of dietary SCars during puberty remodels the developing gut microbiome. Using metagenomic sequencing of fecal samples from mice exposed to high-SCars or control diets, we found that high-SCars exposure profoundly impacted microbial community structure and metabolic potential. Dietary SCars reduced microbial diversity and depleted short-chain fattyacid producing taxa while enriching pathways related to membrane remodeling, branched-chain amino acid biosynthesis, and nucleotide anabolism. These data identify SCars as features of UPFs capable of reprogramming the developing gut microbiota.

## Introduction

There has been a dramatic increase in consumption of ultra-processed foods (UPFs), which are regularly associated with increased chronic disease risk and poorer clinical outcomes[1, 2].In the United States, it is estimated that more than half of daily caloric intake is derived from UPF products. Industrial food processing techniques such as extrusion, retorting, and irradiation introduce and amplify specific chemical modifications in food that, when consumed, may alter key biological processes associated with increased disease risk and complications[3, 4].This may be particularly important during critical developmental windows, such as puberty, when dietary exposures can induce long-lasting biological changes that influence disease susceptibility later in life [5–7]. However, there remains a sparsity of data regarding the specific chemical modifications found in UPFs that influence disease etiology, especially during susceptible periods of biological development.

Advanced glycation end-products (AGEs), advanced lipoxidation end-products (ALEs), and additional environmental carbonyl adducts represent a significant and chemically diverse group of modifications that are amplified in food during manufacturing. Herein grouped under the umbrella term of stable carbonyl adducts (SCars), they represent the final reaction products of spontaneous carbonyl chemistry, in which carbonyl groups on metabolic and oxidative intermediates covalently react with amine groups on proteins, lipids, and nucleic acids [8–11].

UPF consumption represents an ever-increasing source of exogenous SCar exposure [12, 13], as the high heats and pressures applied during cooking and manufacturing accelerate spontaneous carbonyl chemistry within foods, leading to SCars formation and thereby increased exogenous exposure upon consumption [14, 15]. Predominantly characterized within the context of AGEs, both endogenous and exogenous SCars exposure has been linked to chronic metabolic dysfunction and is associated with diabetes, cardiovascular disease, cancer, renal dysfunction, and neurodegenerative disorders in humans and animals [16, 17].

Animal and human studies support SCars *in vivo* bioavailability and tissue accumulation, while dietary intervention studies demonstrate that reducing SCars formation improves metabolic function[16, 18–20]. Epidemiological studies of large human cohorts further associate higher SCars exposure and tissue burden with obesogenic diets, lower socioeconomic status, reduced physical activity, and increased cardiometabolic disease, kidney disease, aging, and cancer outcomes [21–26]. Pre-clinical studies have shown that chronic consumption of SCars-rich diets promotes progressive accumulation in tissues and accelerates metabolic disease, cardiovascular dysfunction, chronic kidney disease, neurodegenerative processes, and cancer progression. Puberty in mice begins around the 25th day of life (post weaning) and represents a critical developmental window during which nutritional exposures can exert long-lasting effects on metabolism and disease susceptibility — analogous to the first 1000 days of human life, when the microbiome is highly malleable and dietary exposures have been linked to food allergy onset, gut permeability, and a wide range of non-communicable disease (NCD) risks [27–29].

Evidence supports that nutritional exposure to SCars has critical implications during this pubertal window, where transient dietary SCars exposure can induce persistent biological changes that influence disease susceptibility long after the initial exposure has ceased [30]. However, the specific role of UPF-associated SCars in those effects remains an understudied area.

Puberty represents a second major window of microbiome plasticity following early life, during which sex hormone shifts, metabolic reorganization, and dietary transitions collectively remodel the gut microbial ecosystem [31]. Microbiomes established during adolescence show reduced resilience compared to adult communities and may be more susceptible to environmental perturbations, including dietary insults [31, 32]. Despite this, the specific effects of dietary chemical modifications on the murine adolescent gut microbiome remain poorly characterized.

The gut microbiome is increasingly recognized as a pivotal mediator linking nutrition to host health, with dietary patterns among the strongest determinants of microbial community structure and function [33–38]. As one of the earliest and most responsive biological systems exposed to dietary SCars [39–44], it represents an important interface for SCars-mediated biological effects. Microbial communities regulate nutrient metabolism, intestinal barrier integrity, immune maturation, bile acid metabolism, and production of bioactive metabolites including SCFAs that influence inflammation, metabolism, and disease risk [45, 46]. Processed food consumption has been associated with lower microbial diversity, reduced abundance of beneficial bacteria, impaired intestinal barrier function, metabolic dysfunction, and disease onset [47–51].

Consequently, alterations in microbial composition and function may represent a novel mechanism through which SCars associated with UPFs influence disease susceptibility. While current literature supports an interaction between dietary SCars and the gut microbiome, their biological importance and causal role in disease pathogenesis remains to be defined.

We therefore sought to determine whether transient dietary SCars exposure is sufficient to remodel the developing gut microbiome by performing shotgun metagenomic sequencing on fecal microbiomes from pubertal mice fed either a high-SCars or control diet. The findings reveal coordinated taxonomic and functional reprogramming of the gut microbiome — characterized by loss of fermentative communities and a shift toward biosynthetic and membrane remodeling processes — with plausible implications for long-term host health.

## Results

### High-SCars Diet Induces Broad Taxonomic and Functional Remodeling of the Gut Microbiome

Differential abundance analysis by MaAsLin3 identified subsets of taxa, enzymes, and metabolic pathways that were differentially abundant between mice fed the high-SCars diet and the control group. At *q* < 0.05, 42 taxonomic features, 81 enzymes, and 15 metabolic pathways were differentially abundant between groups (Table 1). Totals across all FDR thresholds (q < 0.25) reached 96 species genome bins (SGBs), 248 enzyme annotations, and 62 pathways, demonstrating that even a brief, 4-week pubertal exposure to dietary SCars induces broad and coordinated changes in both microbial community structure and metabolic function.

**Table 1.** The FDR-adjusted p-value is listed at different thresholds for SGBs, enzyme commission (EC) numbers, and metabolic pathways, for both abundance and prevalence models. Multiple comparisons were adjusted using the Benjamini–Hochberg procedure (BH), with FDR-adjusted p-values of q < 0.25 or lower considered significant.

| P value (FDR) | # SGBs (abundance) | # SGBs (prevalence) | # ECs (abundance) | # ECs (prevalence) | # Pathways (abundance) | # Pathways (prevalence) |
| --- | --- | --- | --- | --- | --- | --- |
| 0.10–0.25 | 20 | 93 | 79 | 0 | 17 | 0 |
| 0.05–0.10 | 16 | 0 | 36 | 0 | 14 | 0 |
| 0.01–0.05 | 42 | 0 | 81 | 3 | 15 | 0 |
| <0.01 | 18 | 0 | 52 | 0 | 16 | 0 |
| **Totals** | **96** | **93** | **248** | **3** | **62** | **0** |

### High-SCars Exposure Reduces Gut Microbial Diversity and Reshapes Community Composition

Alpha diversity analysis revealed that pubertal exposure to high-SCars significantly decreased microbial diversity relative to regular diet (Figure 1A). Chao1 richness (*P* < 0.003), Shannon diversity (*P* < 0.001), and Simpson diversity (*P* < 0.003) were all significantly decreased in the high-SCars group, indicative of lower microbial richness and evenness. Beta diversity analysis likewise showed substantial differences in overall community composition. Bray–Curtis principal coordinate analysis (PCoA) distinctly separated high-SCars from regular diet communities, with diet explaining 41% of total community variation (PERMANOVA R² = 0.41, *P* < 0.001) (Figure 1B). The first two PCoA axes captured 48.3% and 19% of total community variation, respectively, with high-SCars and regular diet samples forming visually non-overlapping clusters. Notably, high-SCars communities exhibited greater within-group dispersion than regular diet communities (Figure 1B), suggesting that SCars exposure produces heterogeneous microbial responses rather than a single stereotyped disruption. Together, these analyses demonstrate that transient exposure to dietary SCars during puberty dramatically reshapes the gut microbiota.

**Figure 1.**
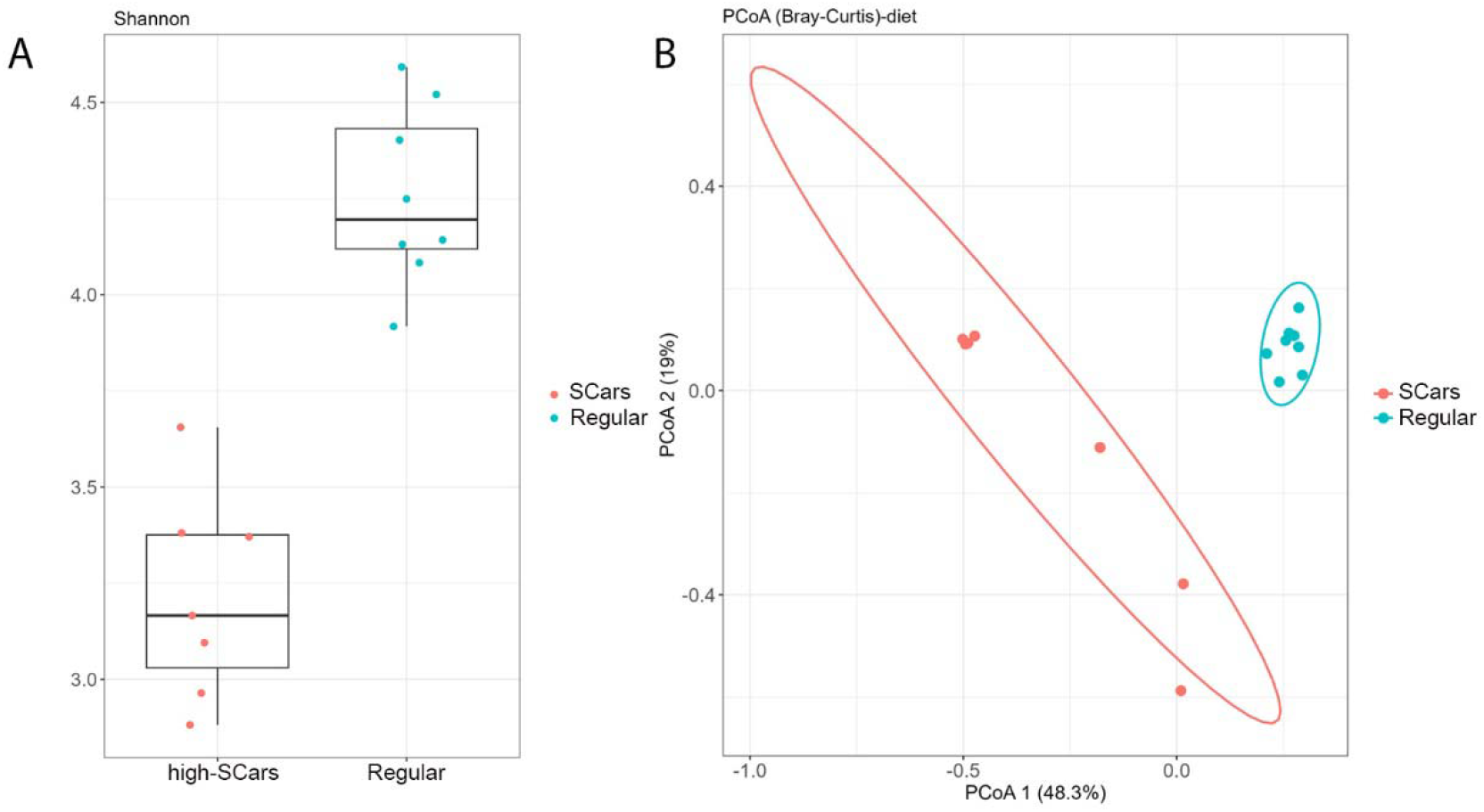
Alpha and beta diversity of gut microbiota from high-SCars and Regular diet groups. (A) Shannon diversity index for microbiota from mice fed high-SCars or regular diets, showing significantly reduced species richness and evenness in the high-SCars group. (B) Bray–Curtis PCoA demonstrating community-level separation by diet. Each point represents one animal; ellipses denote 95% confidence regions. Diet explained 41% of total community variation (PERMANOVA R² = 0.41, *P* < 0.001).

**Figure 2.**
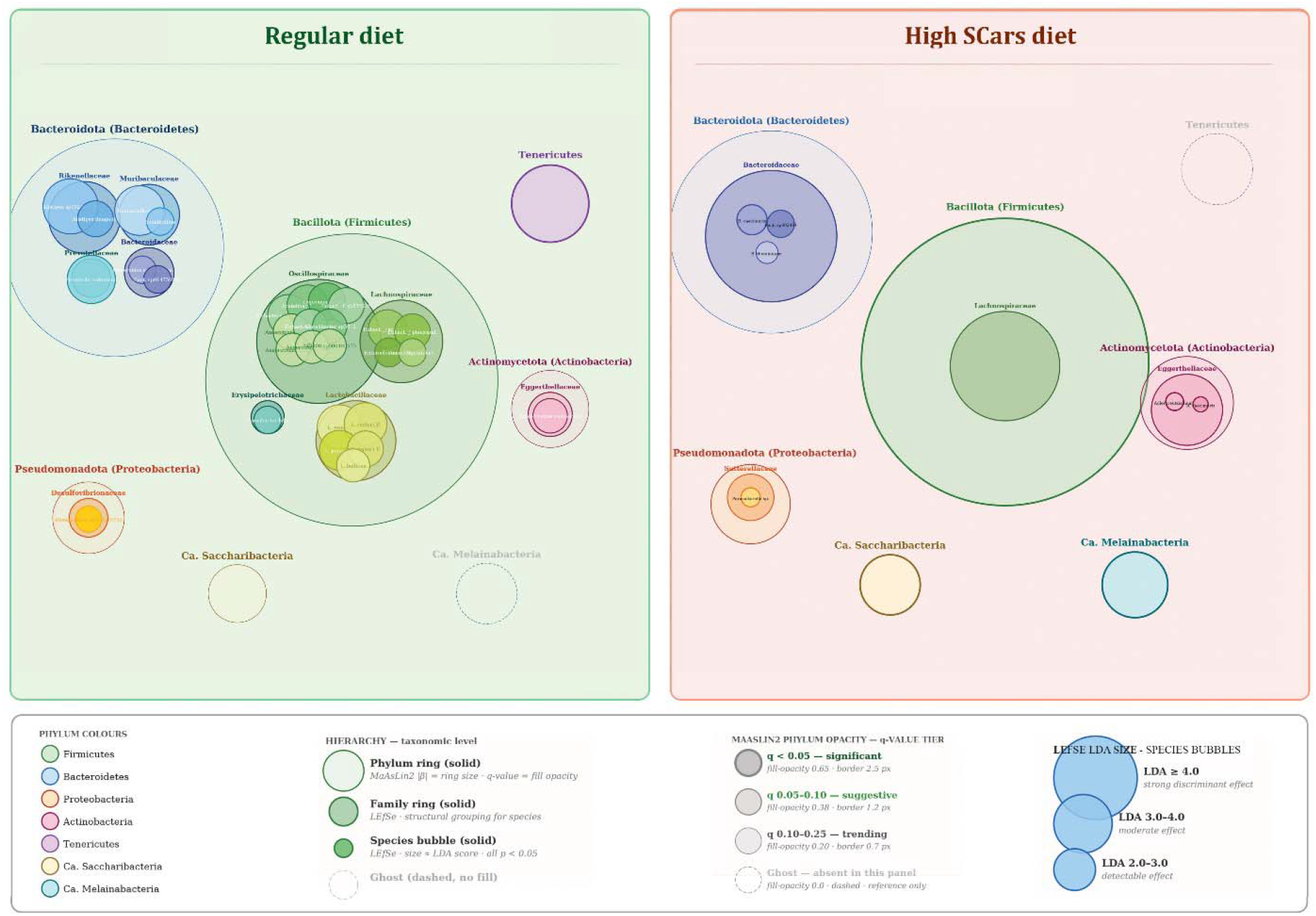
Differentially abundant bacterial taxa associated with regular and high-SCars diets. Shotgun metagenomic profiling identified diet-associated microbial taxa using LEfSe and MaAsLin3 analyses. Taxa are organized hierarchically by phylum, family, and species, with bubble size proportional to LEfSe effect size (LDA score). Relative to the regular diet, pubertal high-SCars exposure resulted in reduced taxonomic complexity, and selective enrichment of candidate phyla *Ca.* Saccharibacteria and *Ca.* Melainabacteria, whereas the regular diet was associated with enrichment of numerous taxa within the Bacteroidota and Bacillota phyla.

Shotgun metagenomic profiling identified 338 species distributed across seven bacterial phyla. More than 80 taxa exhibited differential abundance as a function of diet (q < 0.25; Table 1), demonstrating broad-scale taxonomic remodeling following pubertal high-SCars consumption. The regular diet supported nearly six-fold more enriched taxa than the high-SCars diet (29 versus 5 species), indicating a marked contraction of microbial diversity and taxonomic complexity. Taxa enriched under each dietary condition were primarily members of the two most abundant phyla, Bacillota (Firmicutes) and Bacteroidota (Bacteroidetes).

Beyond the dominant phyla, high-SCars diet was also associated with a marked expansion of Ca. Melainabacteria (SGB44380), detected in 6 of 7 high-SCars samples at substantially higher relative abundance (mean 1.21%) compared to 4 of 8 regular diet samples at near-trace levels (mean 0.07%). These ultra-small, obligate episymbiotic organisms represent a poorly characterized component of the gut microbiome, and their selective enrichment under dietary carbonyl stress is a novel observation warranting further investigation. In contrast, Ca.

Saccharibacteria was detected across all regular diet samples but showed reduced prevalence in high-SCars microbiomes (3/7 samples), with two high-SCars outlier samples driving a higher group mean — a pattern more consistent with community disruption than directional enrichment. Concurrently, a Proteobacteria (Pseudomonadota) representative was enriched in high-SCars microbiomes (q = 0.086), consistent with Proteobacteria bloom patterns commonly reported in states of gut dysbiosis and mucosal inflammation. In contrast, microbial taxa within Mycoplasmatota (previously Tenericutes; MaAsLin3 coef +3.94, q = 0.024), as well as abundant taxa within the Lachnospiraceae, Oscillospiraceae, Rikenellaceae, and Prevotellaceae families, were selectively enriched by regular dietary consumption.

### High-SCars Diet Selectively Remodels the Bacteroidota Phylum

Taxonomic analysis revealed Bacteroidota as the phylum most differentially affected by pubertal high-SCars diet (Figure 3A). While regular diet promoted enrichment of seven species from four families (Rikenellaceae, Muribaculaceae, Prevotellaceae, and Bacteroidaceae), high-SCars diet was associated with decreased taxonomic breadth, enriching only three species from the single family Bacteroidaceae. MaAsLin3 differential abundance analysis validated enrichment of *Bacteroides caecimuris* (FDR 1.44e-02; Coef. 4.95), *Bacteroides thetaiotaomicron* (FDR 1.517e-02; Coef. 1.48), and *Bacteroides* sp002491635 (FDR 1.498e-02; Coef. 2.10) in high-SCars microbiomes (Figure 3B). Notably, the families enriched under regular diet conditions — Rikenellaceae, Muribaculaceae, and Prevotellaceae — are broadly associated with dietary fiber fermentation and maintenance of gut homeostasis, while the Bacteroidaceae species enriched under high-SCars conditions, particularly *B. thetaiotaomicron*, are known host-glycan foragers that expand when mucosal barrier function is compromised. This suggests a functional shift in Bacteroidota community behavior — from fiber fermentation to opportunistic host-glycan utilization — rather than simple taxonomic replacement. Taken together, pubertal exposure to dietary SCars differentially restructures Bacteroidota by narrowing taxonomic diversity and selectively enriching host-adapted Bacteroides species.

**Figure 3.**
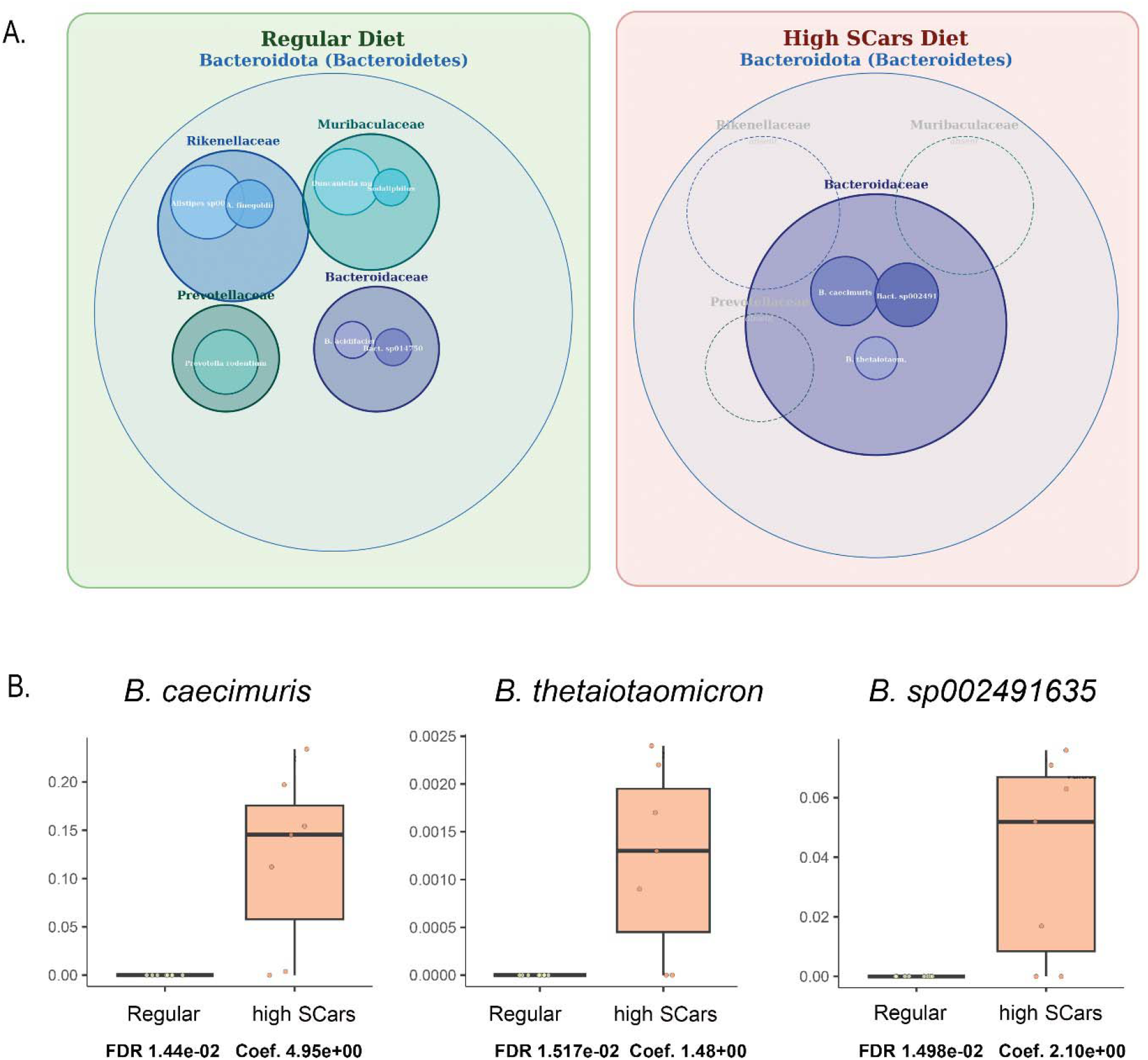
Bacteroidota taxa differentially regulated by SCars diet exposure. (A) Hierarchical bubble plot showing differentially abundant Bacteroidota taxa, organized by family (solid ring) and species (filled bubbles), with bubble size proportional to LEfSe LDA score. Regular diet microbiomes were enriched for species across four Bacteroidota families, while high-SCars microbiomes showed enrichment restricted to three species within Bacteroidaceae. (B) MaAsLin3 differential abundance coefficients for *Bacteroides* species enriched in high-SCars microbiomes, shown with FDR-adjusted q-values and effect size coefficients.

### High-SCars Diet Depletes SCFA-Producing Bacillota

Within the Bacillota phylum, a greater diversity of species spanning four families — Oscillospiraceae, Lactobacillaceae, Lachnospiraceae, and Erysipelotrichaceae was observed contrasted to less significant enriched taxa in SCars diet (Figure 4A). Many taxa that were reduced in SCars diet samples were recognized producers of SCFAs, specifically butyrate. The depletion of a butyrogenic guild encompassing taxa from Lachnospiraceae (*P* = 0.0205), Oscillospiraceae, and Odoribacteraceae (*P* = 0.0037) is shown in Figure 4B, and C. Among the most depleted taxa were *Oscillibacter* species, *Acetatifactor muris*, and *Neglectibacter* sp., all members of the butyrogenic guild. Because establishment of SCFA-producing communities is important for immune and epithelial maturation during adolescence [32], depletion of these organisms during puberty may have broader consequences than similar perturbations occurring later in adulthood.

**Figure 4.**
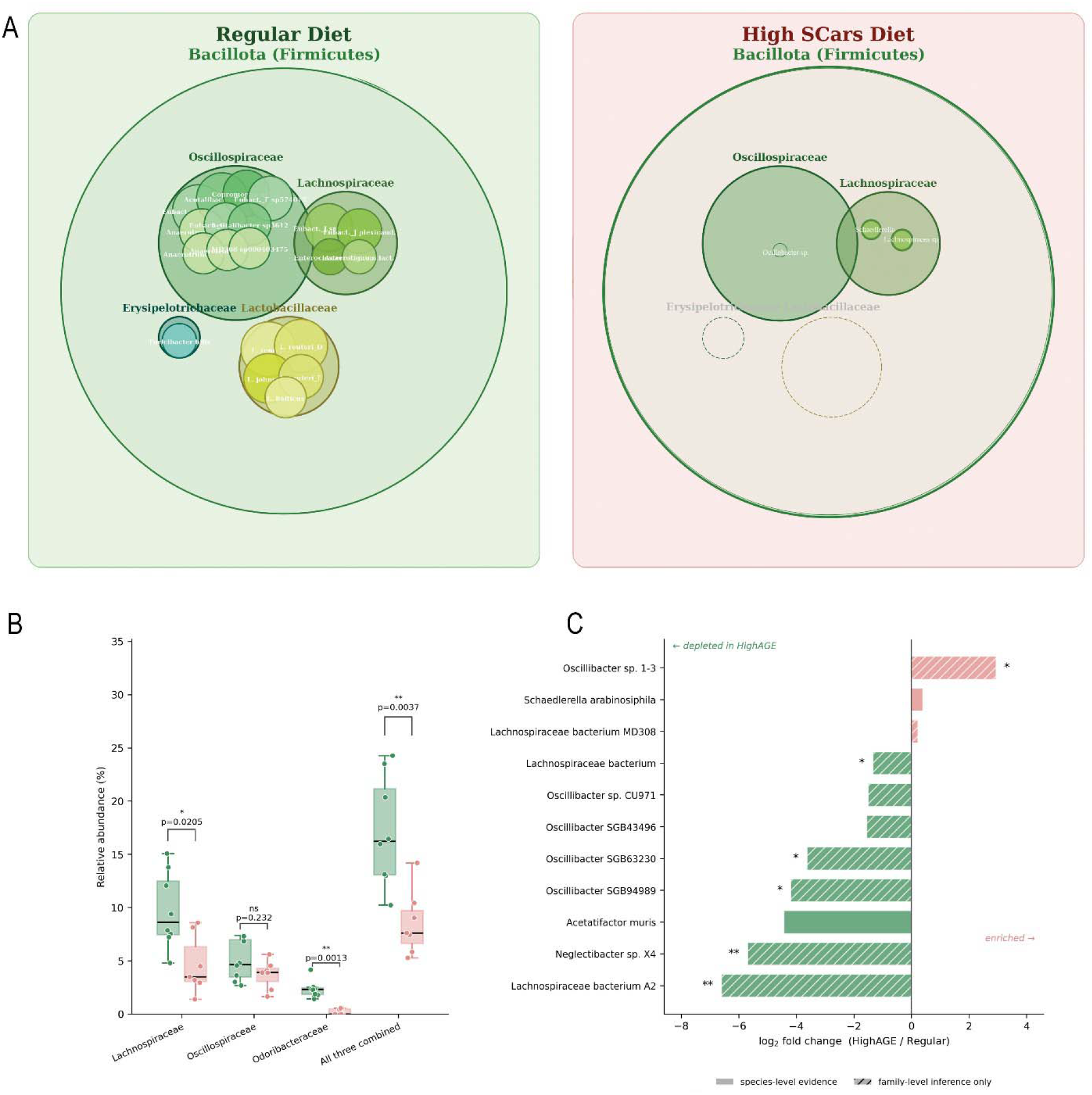
Pubertal high-SCars exposure causes marked depletion of SCFA-producing Firmicutes (Bacillota). (A) Hierarchical taxonomic representation of differentially abundant Bacillota taxa identified by LEfSe and MaAsLin3. Taxa are organized by phylum, family, and species, with bubble size proportional to LEfSe LDA score. Compared with the regular diet, taxonomic diversity was greatly reduced in the high-SCars diet. (B) Differential abundance of a subset of taxa with known association with butyrate production. (C) Log fold-change of candidate butyrate producers at family level, showing depletion of Lachnospiraceae, Oscillospiraceae, and Odoribacteraceae combined under high-SCars conditions.

### Pubertal High-SCars Exposure Remodels Functional Metabolic Pathways

Univariate functional pathway analysis using HUMAnN revealed 62 differentially abundant metabolic pathways between dietary groups (q < 0.25), 31 of which remained significant under stricter correction (q < 0.05; Table 1). Distinct metabolic programs were associated with each diet (Figures 5, 6). High-SCars microbiomes were enriched for pathways involved in branched-chain amino acid (BCAA) biosynthesis (BCAA superpathway, L-isoleucine biosynthesis I and III), nucleotide anabolism (pyrimidine biosynthesis II, guanosine nucleotides degradation III), NAD salvage (PNC IV cycle), lipid and membrane biosynthesis (phospholipid biosynthesis I, CMP-pseudaminate biosynthesis, UDP-N-acetylglucosamine, CMP-Kdo biosynthesis), and vitamin B6 biosynthesis (pyridoxal 5’-phosphate biosynthesis I and salvage), consistent with increased anabolic and membrane remodeling functions. In contrast, microbiomes from mice consuming the regular diet preferentially enriched pathways associated with fermentation and SCFA production (glycolysis IV, pyruvate fermentation to butanoate, *Clostridium acetobutylicum* acidogenic fermentation superpathway), sugar acid degradation (D-galacturonate, D-fructuronate, glucuronides, and stachyose degradation), amino acid catabolism (L-histidine degradation I), and TCA cycle metabolism.

**Figure 5.**
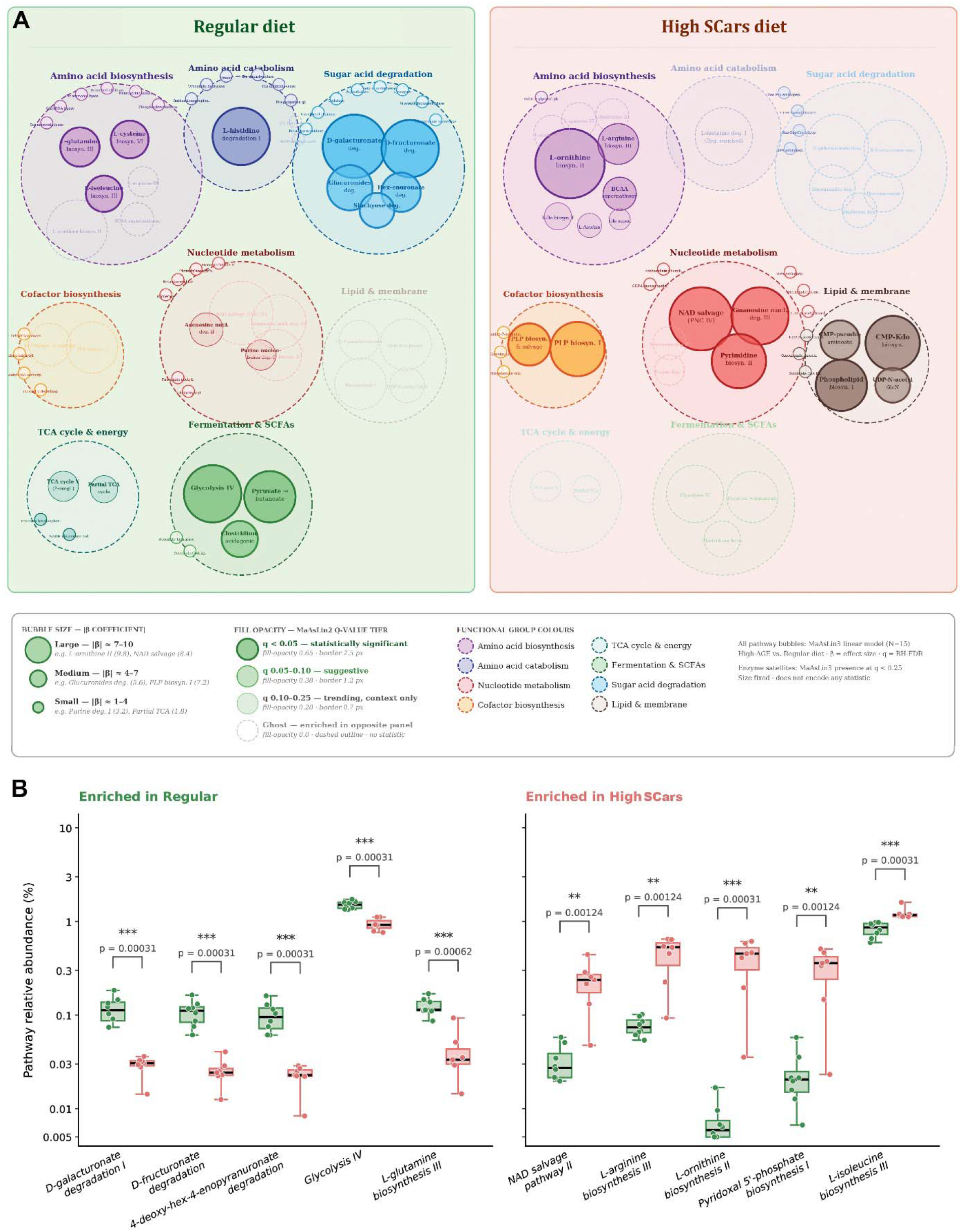
Differentially abundant microbial metabolic pathways following pubertal high-SCars exposure. **5A.** Functional pathways were identified using HUMAnN and differential abundance determined using MaAsLin3. Pathways are grouped into major functional categories and displayed as a dual-panel bubble plot (regular diet, left; high-SCars diet, right). Bubble size reflects the MaAsLin3 |β coefficient|; fill opacity reflects q-value tier (q < 0.05 = statistically significant; q 0.05–0.10 = suggestive; q 0.10–0.25 = trending). Ghost bubbles (dashed outline) indicate pathways enriched in the opposite panel. Green and red indicate enrichment in regular and high-SCars microbiomes, respectively. **5B.** Community level HUMAnN pathway abundances, TSS normalized (% of total mapped pathway abundance); log10 y-axis. Boxes span the IQR with median and full range; points are individual mice. Two-sided Mann-Whitney U; *** p < 0.001, ** p < 0.01. p= 0.00031 is the smallest attainable with n = 8 vs. 7. Pathways are the five most significant in Regular and High SCars enriched pathways.

**Figure 6.**
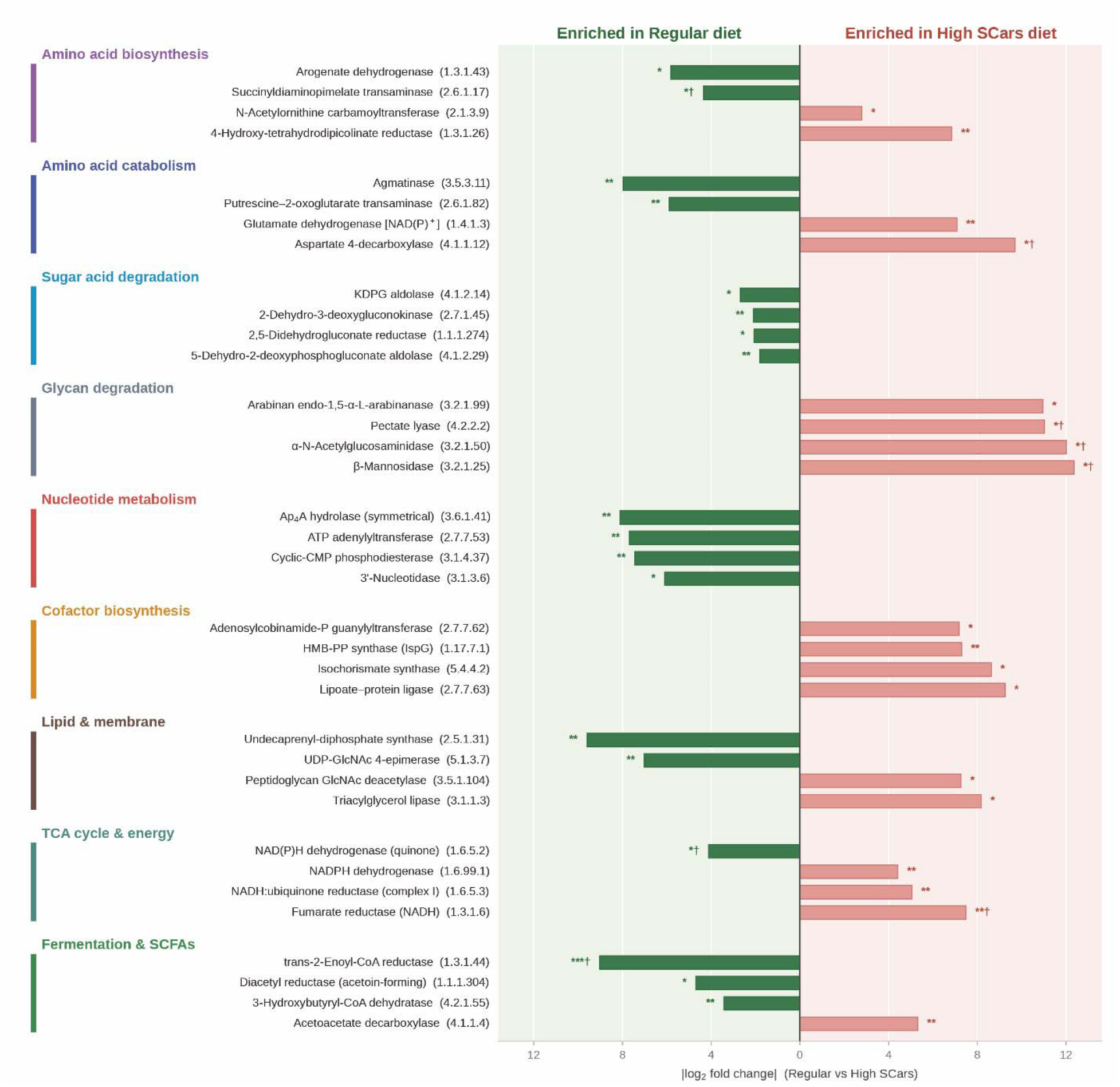
Differential Enrichment of Enzymes between Regular and High-SCars Diets. The four largest effects per functional group among the 436 ECS significant at q < 0.05 (*** q < 0.001, ** q< 0.01, * q< 0.05).

A particularly notable observation was the directional switch in nucleotide metabolism between dietary conditions. Regular diet microbiomes were enriched for nucleotide catabolic pathways (adenosine nucleotide degradation II), while high-SCars microbiomes were enriched for nucleotide biosynthetic pathways (pyrimidine biosynthesis II, guanosine nucleotides degradation III, NAD salvage pathway PNC IV cycle). This shift from nucleotide recycling to de novo synthesis is consistent with a community-level transition toward active microbial expansion or stress response. Similarly, the sugar acid degradation cluster — encompassing D-galacturonate, D-fructuronate, glucuronides, and stachyose degradation — was selectively enriched in regular diet microbiomes, reflecting the capacity of these communities to ferment complex plant-derived polysaccharides, a function essentially absent in high-SCars microbiomes.

Among all differentially abundant pathways, L-ornithine biosynthesis II showed the largest effect size (coef −5.7, q = 0.001), being strongly enriched in high-SCars microbiomes. L-ornithine is the biosynthetic precursor to polyamines including putrescine and spermidine, which regulate microbial stress responses and host-microbe interactions; its enrichment may reflect a stress-adaptive biosynthetic response to dietary carbonyl exposure and warrants further investigation given the known roles of microbially derived polyamines in host epithelial biology.

### Pubertal High-SCars Exposure Remodels the Functional Enzyme Repertoire

Functional metagenomic profiling identified 1,659 unique enzyme commission (EC) numbers, 248 of which were significantly different between dietary groups (q < 0.25), with 81 remaining significant under strict correction (q < 0.05; Table 1). Regular diet-fed mice had microbiomes enriched for enzymes associated with SCFA fermentation, central carbon metabolism, TCA cycle function, amino acid catabolism, and histidine degradation. High-SCars microbiomes preferentially enriched carbohydrate-active enzymes (CAZymes), nucleotide metabolism enzymes, amino acid biosynthetic enzymes, cofactor biosynthesis, and lipid and membrane remodeling features consistent with a shift toward nutrient scavenging and membrane restructuring.

Among the enzymes enriched in high-SCars microbiomes, several CAZymes — including β-galactosidase, α-galactosidase, β-glucosidase, β-glucuronidase, cellulase, xylan 1,4-β-xylosidase, and related glycosidases were significantly increased. The enrichment of CAZymes in high-SCars microbiomes despite concurrent loss of canonical SCFA-producing taxa presents an apparent paradox. Rather than reflecting enhanced dietary fiber fermentation, CAZyme enrichment in this context most likely represents a shift toward opportunistic host-glycan foraging — a metabolic strategy employed by *Bacteroides* species under conditions of mucosal barrier disruption. This interpretation is supported by the concurrent enrichment of *B. thetaiotaomicron* and *B. caecimuris*, both well-characterized host-glycan specialists[52]. Regular diet microbiomes were also enriched for peptide processing enzymes including rhomboid protease and multiple aminopeptidases, suggesting greater proteolytic capacity and amino acid cycling consistent with a more metabolically diverse community.

## Discussion

This study identifies SCars as an underappreciated component of ultra-processed foods capable of remodeling the developing gut microbiome during puberty (Figure 7). Transient exposure to a high-SCars diet induced coordinated taxonomic, enzymatic, and metabolic reprogramming characterized by reduced microbial diversity, depletion of health-promoting SCFA-producing bacteria, enrichment of biosynthetic and proteolytic functions, and a shift away from saccharolytic fermentation. The developing microbiome is substantially more plastic during puberty than in adulthood, and transient dietary exposures during this period may produce biological changes that persist well beyond the exposure itself [30]. Our findings complement previous observations that pubertal SCars exposure induces persistent mammary gland remodeling [53] and extend this concept to demonstrate that the developing gut microbiome is susceptible to dietary carbonyl stress. Because puberty represents a critical developmental window during which dietary exposures may have lasting biological consequences, these findings identify dietary SCars as a plausible mechanistic link between ultra-processed food consumption and chronic disease risk.

**Figure 7.**
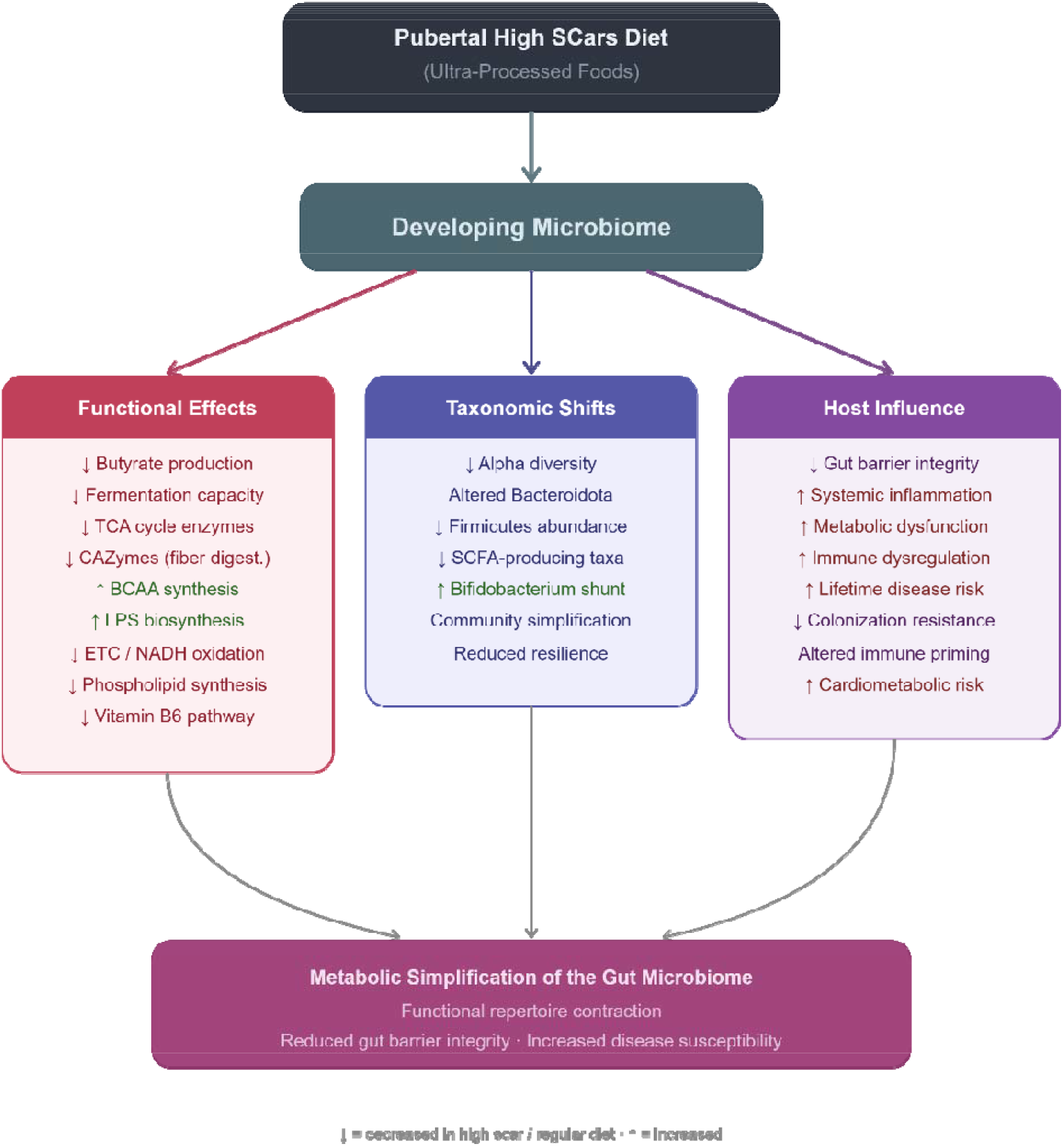
Proposed model illustrating how transient pubertal exposure to a high-SCars diet remodels the gut microbiome and its metabolic potential. Pubertal high-SCars diet exposure drives coordinated taxonomic simplification and functional reprogramming of the gut microbiome, with downstream implications for host barrier integrity, immune maturation, and disease susceptibility.

High-SCars consumption during puberty caused significant reductions in alpha diversity and shifted overall microbiome community structure. This observation is consistent with prior reports showing that SCars are associated with restructuring of the microbial ecosystem and metabolic disease [39–44]. Loss of microbial diversity per se is not necessarily pathological; however, the concurrent depletion of several health-promoting taxa alongside enrichment of alternative metabolic pathways suggests that dietary SCars drive a functional reorganization of the developing gut microbial ecosystem — one likely to influence health consequences well beyond the period of dietary exposure. Notably, high-SCars communities exhibited greater within-group dispersion than regular diet communities, suggesting that SCars exposure produces individualized, heterogeneous microbial disruption rather than a single stereotyped response, consistent with the known influence of host genetics and prior microbial history on resilience to dietary perturbation.

Among the most novel taxonomic observations was the enriched detection of candidate phyla radiation (CPR) lineage -*Ca.* Melainabacteria in high-SCars microbiomes, with only trace representatives detected in any regular diet sample. CPR organisms are ultra-small, obligate epibionts with highly reduced metabolic capabilities; their enrichment under high dietary carbonyl stress may reflect ecological opportunities created by community restructuring, or alternatively may serve as a marker of microbiome destabilization. While the functional significance of these organisms in mammalian gut ecosystems remains largely unknown, their selective appearance in a dysbiotic dietary context warrants further investigation.

Dietary exposure to SCars during puberty was strongly associated with depletion of multiple SCFA-producing Bacillota as well as pathways responsible for butyrate fermentation. Butyrate-producing bacteria play key roles in colonocyte metabolism, epithelial integrity, and immunomodulation, and have been implicated in disease states ranging from inflammatory bowel disease to colorectal cancer and neurodegeneration [54, 55]. Because establishment of SCFA-producing communities is important for immune and epithelial maturation during adolescence [32], depletion of these organisms during puberty may have broader implications than similar perturbations occurring later in adulthood. Our data therefore suggest that consumption of SCars may promote loss of microbial functions that support intestinal homeostasis during a particularly sensitive developmental period.

Microbiomes from high-SCars diet mice underwent profound metabolic reprogramming. Enrichment of pathways involved in BCAA biosynthesis, nucleotide anabolism, NAD salvage, lipid and membrane biosynthesis, and carbohydrate-active enzymes coincided with depletion of pathways related to glycolysis, TCA cycle metabolism, fermentation, and butyrate production.

These findings suggest that exposure to dietary SCars promotes a transition away from a metabolically cooperative community that ferments dietary fiber toward one centered on biosynthetic processes, host glycan utilization, and nutrient scavenging — a metabolic signature consistent with a community under environmental stress [56].

A particularly notable observation was the directional switch in nucleotide metabolism: regular diet communities were enriched for nucleotide catabolic pathways (adenosine nucleotide recycling), while high-SCars communities were enriched for nucleotide biosynthetic pathways (pyrimidine biosynthesis, NAD salvage, guanosine metabolism). This shift from recycling to de novo synthesis is consistent with a proliferating or stress-adapted microbial population responding to carbonyl challenge and may also contribute to altered host NAD homeostasis given the known exchange of NAD precursors between gut bacteria and the host.

The strong enrichment of L-ornithine biosynthesis II — the pathway with the largest effect size among all differentially abundant pathways (coef − 5.7, q = 0.001) — in high-SCars microbiomes raises the possibility that polyamine production is elevated in these communities. Microbially derived polyamines, including putrescine and spermidine, regulate host epithelial proliferation and immune function; their dysregulation has been implicated in colorectal carcinogenesis, providing a potential mechanistic link between SCars-induced microbiome remodeling and cancer risk that warrants experimental validation.

The concurrent enrichment of CAZymes alongside depletion of fiber-fermenting taxa presents an interpretive paradox. Rather than reflecting enhanced dietary fiber fermentation, CAZyme enrichment in high-SCars microbiomes most likely represents a shift toward host-derived glycan foraging — a metabolic strategy employed by opportunistic *Bacteroides* species under conditions of mucosal barrier disruption. This interpretation is supported by the selective enrichment of *B. thetaiotaomicron* and *B. caecimuris*, both well-characterized host-glycan specialists that expand when mucosal integrity is compromised [52].

We also observed enrichment of lipopolysaccharide biosynthesis, peptidoglycan metabolism, and membrane remodeling pathways in high-SCars microbiomes, changes consistent with processes thought to drive metabolic endotoxemia, intestinal leak, and chronic inflammation. While our metagenomic approach cannot quantify endotoxin production or direct host exposure, these pathway-level changes are consistent with a pro-inflammatory microbial phenotype.

Experimental validation including metabolomic profiling, endotoxin quantification, and host immune measurements will be required to confirm these hypotheses and determine whether these functional alterations influence maturation of host immune and metabolic pathways during adolescence.

There are several important limitations to acknowledge. The duration of exposure was brief, sample size was limited, and rodent models do not necessarily translate directly to human health outcomes. Functional alterations were inferred from metagenomic sequencing data and were not confirmed by metabolomic profiling or direct physiological measurements of SCFAs, circulating BCAAs, intestinal permeability, or inflammatory markers. Our use of autoclaving as a model of food processing increases SCars formation but also alters food texture, digestibility, and potentially other bioactive compounds — a confound that should be acknowledged when interpreting results. Microbiome composition was characterized at a single time point immediately following pubertal exposure; longitudinal studies will be required to establish whether these alterations persist into adulthood or are reversed following dietary normalization.

Ultra-processed foods comprise a chemically diverse class of products for which the mechanisms underlying adverse health effects remain incompletely understood. Our findings support the concept that processing-derived stable carbonyl adducts represent a previously underrecognized chemical feature of these foods capable of reprogramming the developing gut microbiome during puberty. By identifying coordinated taxonomic and functional remodeling following transient pubertal SCars exposure, this study provides a plausible mechanistic framework linking industrial food processing, microbiome dysfunction, and chronic disease risk. Critically, dietary SCars are a potentially modifiable exposure: food processing methods that minimize Maillard reaction chemistry — lower cooking temperatures, shorter processing times, moisture retention — measurably reduce SCars content without compromising food safety. If microbiome reprogramming during adolescence represents a mechanistic pathway to adult disease, reducing SCars exposure during this developmental window may constitute a tractable and impactful dietary intervention.

## Methods

### Mouse Caging and Diet Preparation

Thirty FVB/N mice (JAX stock #001800) were caged (15 cages) and divided into two groups: ‘Regular’ (n = 8) and ‘High SCars’ (n = 7). Regular diet mice received TestDiet 58G7 (TestDiet, Richmond, IN); high-SCars mice received the same diet autoclaved at 120°C for 15 min, following a published method [21]. Mice had *ad libitum* access to water throughout the 4-week dietary exposure period.

### AGE/SCars Measurement

Concentrations of carboxymethyl lysine (CML) and methylglyoxal-derived hydroimidazolone (MG-H1) in mouse diets were determined by enzyme-linked immunosorbent assay as described previously[53], as quantitative indicators of AGE/SCars levels across both diets (Supplementary Figure 1).

### DNA Extraction

Stool pellet samples were frozen at −20°C until DNA extraction using the QIAamp PowerFecal Pro DNA Kit (Qiagen) according to the manufacturer’s specifications.

### Library Preparation and Sequencing

DNA concentrations were measured by fluorometric quantification (Qubit, Thermo Fisher Scientific) and libraries were prepared using the Illumina DNA Prep kit according to the manufacturer’s specifications. Libraries were normalized and sequenced on a NextSeq2000 using a P3 cartridge. After initial quality screening, all samples had a minimum of 120 million reads for downstream analyses.

### Bioinformatic Analyses

The KneadData pipeline was used to trim reads and remove host contamination using a C57BL/6J reference genome (GRCm39). Quality-controlled reads were processed with the bioBakery pipeline (MetaPhlAn 4.0.6, ChocoPhlAn database: Oct22) to generate taxonomic profiles, and with HUMAnN v3.7 to generate functional profiles. Quality control and statistical analyses (alpha diversity, beta diversity, linear modeling) were performed in R v4.4.3 using the following packages: phyloseq, vegan, rstatix, adonis2, ape, and MaAsLin3. MaAsLin3 (61) was used for linear and logistic regression to associate abundance and prevalence of taxonomic, functional, and enzyme features with dietary group, correcting for sequencing depth (Supplementary Figures 2, 3, and 4 respectively). The taxonomic table was filtered for features with greater than 0.1% relative abundance in more than 10% of samples. Multiple comparisons were adjusted using the Benjamini–Hochberg procedure (BH), with FDR-adjusted p-values of q ≤ 0.25 reported as significant.

### Data Visualization

Visualizations of bioBakery outputs (MaAsLin3, LEfSe, MetaPhlAn) were created using R v4.4.3. Generative AI tools (Claude, Anthropic) were used to generate Python code for hierarchical bubble visualizations (Figures 2–5) using MaAsLin3 and LEfSe outputs and MetaPhlAn taxonomic outputs were also used with Claude generated python code to visualize inferred hierarchical relationships in circular cladograms (Supplementary Figure 5).

### Use of Generative AI

The generative AI tool Claude (Anthropic) was used during the preparation of this manuscript to assist with bioinformatic code generation, data visualization, and editorial proofreading. All scientific content, analyses, interpretations, and conclusions were performed and verified by the authors, who reviewed and assume full responsibility for the integrity of the final manuscript.

## Acknowledgements

We would like to acknowledge the support of the Office of Applied Science at the Center for Veterinary Medicine of the U.S. Food and Drug Administration for support of this work.

## Author Credit Roles

VJF, DPT, AO conceived of the study. JL and VJF managed mice studies, BK and AO coordinated molecular laboratory work. AO and BK conducted bioinformatic analyses. AO, VJF, and DPT wrote the paper.

## Ethics Approval and Consent to Participate

Animal work was approved by the Institutional Animal Care and Use Committee of Virginia Commonwealth University and conducted in accordance with institutional and local requirements.

## Data Availability Statement

All data have been submitted to NCBI under BioProject PRJNA1522370 https://www.ncbi.nlm.nih.gov/search/all/?term=PRJNA1522370

## Supplementary Figures

**Supplementary Figure 1.**
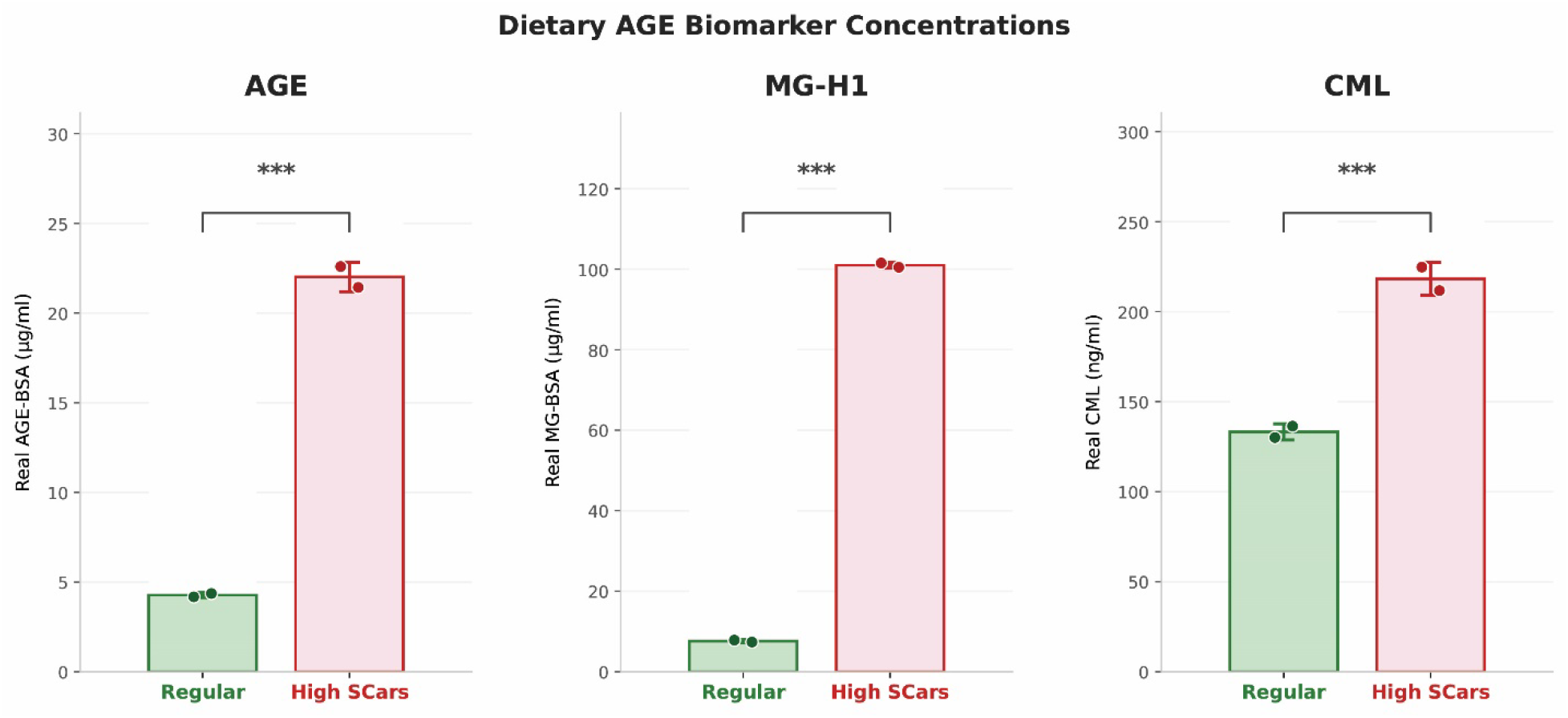
SCars Levels. Concentrations of carboxymethyl lysine (CML) and methylglyoxal-derived hydroimidazolone (MG-H1) in mouse diets were determined by enzyme-linked immunosorbent to estimate AGE/SCars levels as described previously[53].

**Supplementary Figure 2.**
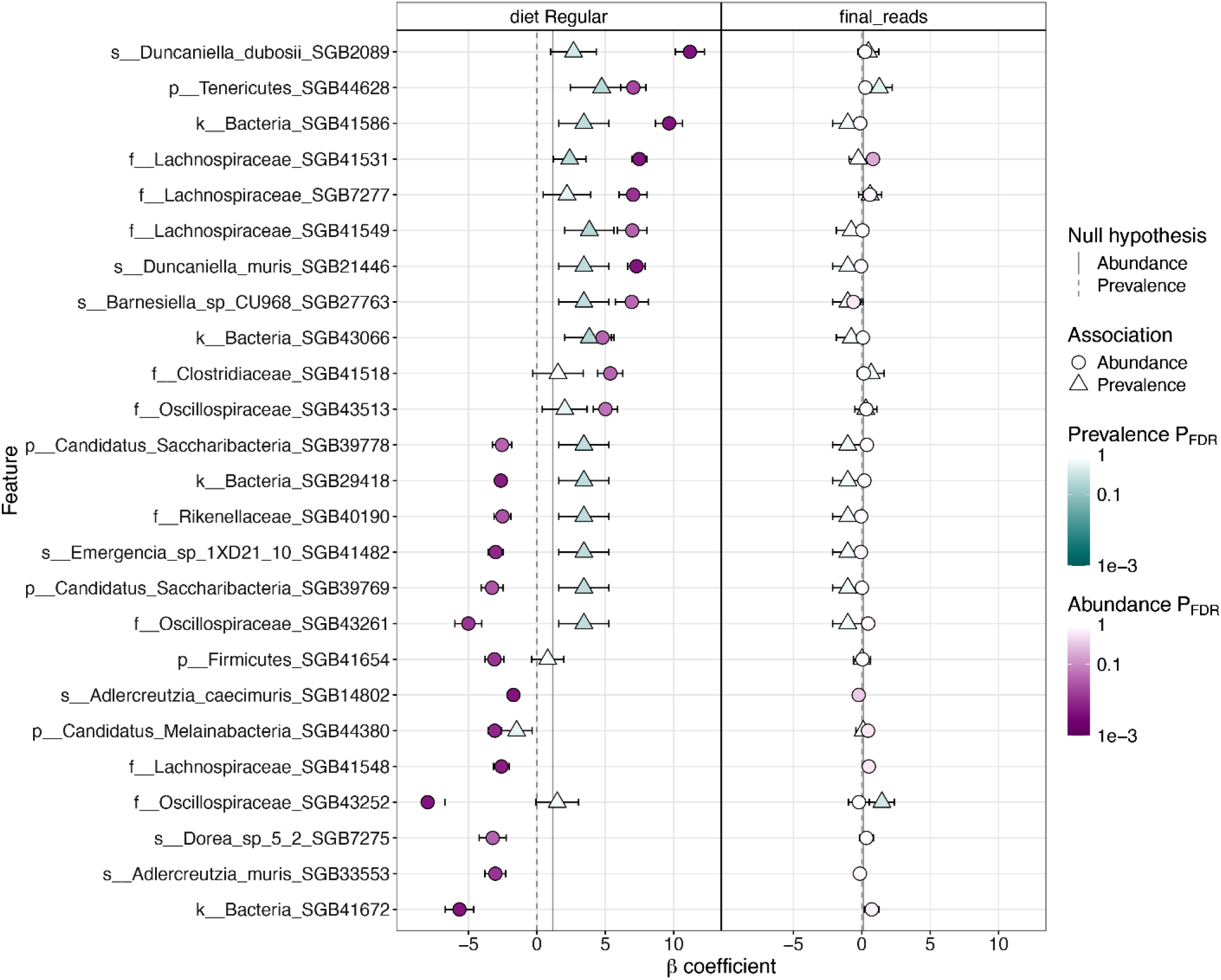
Microbial taxonomy associated with diet, identified with MaAsLin 3. Coefficient plot for the 25 taxonomic signals (profiled with MetaPhlAn 4.0.6, ChocoPhlAn database: Oct22) with the strongest diet associations in a multivariable MaAsLin 3 model. The model included diet ([Regular vs. reference diet]) and sequencing depth (final_reads). Each row is one pathway, ordered by its diet coefficient.

**Supplementary Figure 3.**
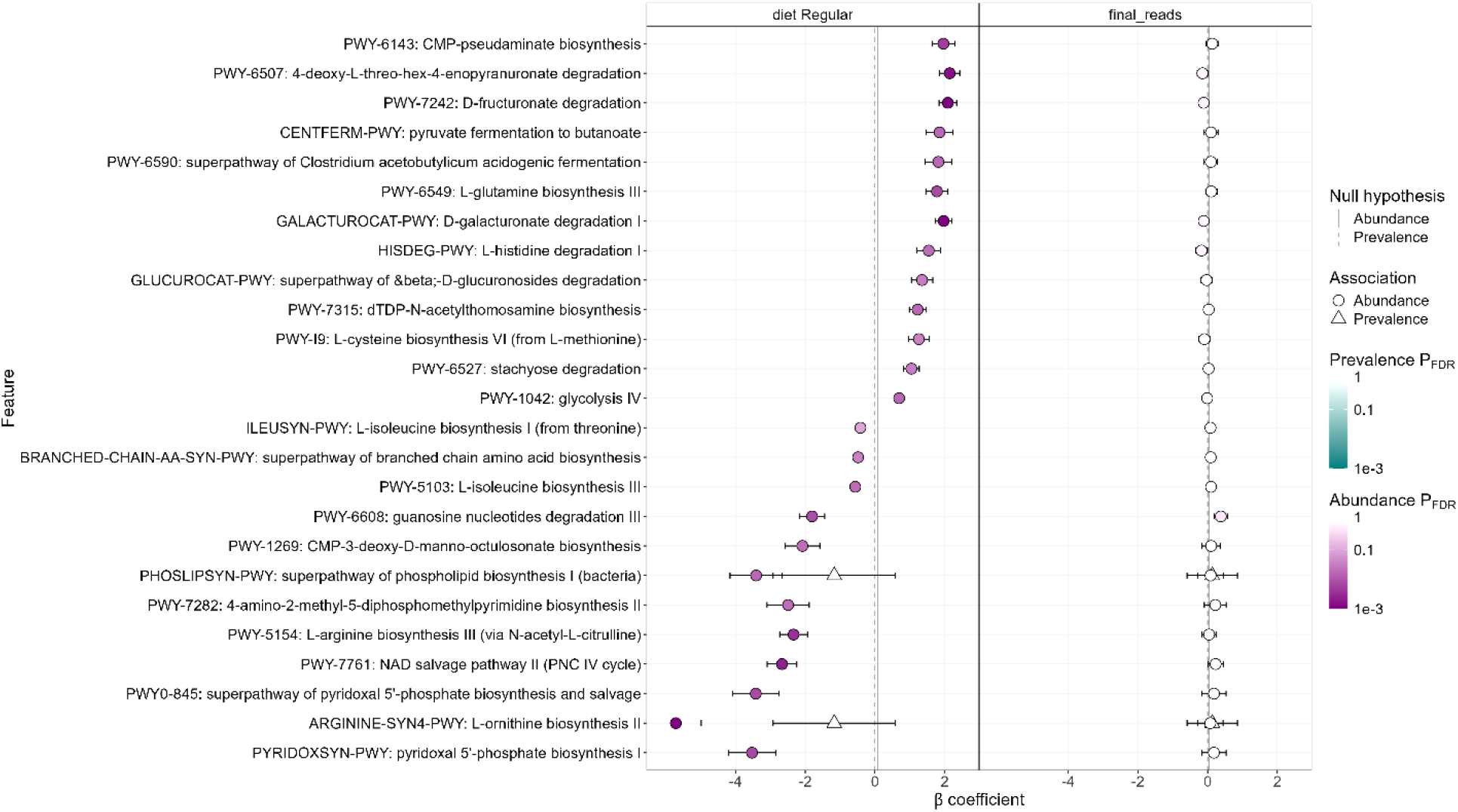
Microbial metabolic pathways associated with diet, identified with MaAsLin 3. Coefficient plot for the 25 MetaCyc pathways (profiled with [HUMAnN 3]) with the strongest diet associations in a multivariable MaAsLin 3 model. The model included diet ([Regular vs. reference diet]) and sequencing depth (final_reads). Each row is one pathway, ordered by its diet coefficient.

**Supplementary Figure 4.**
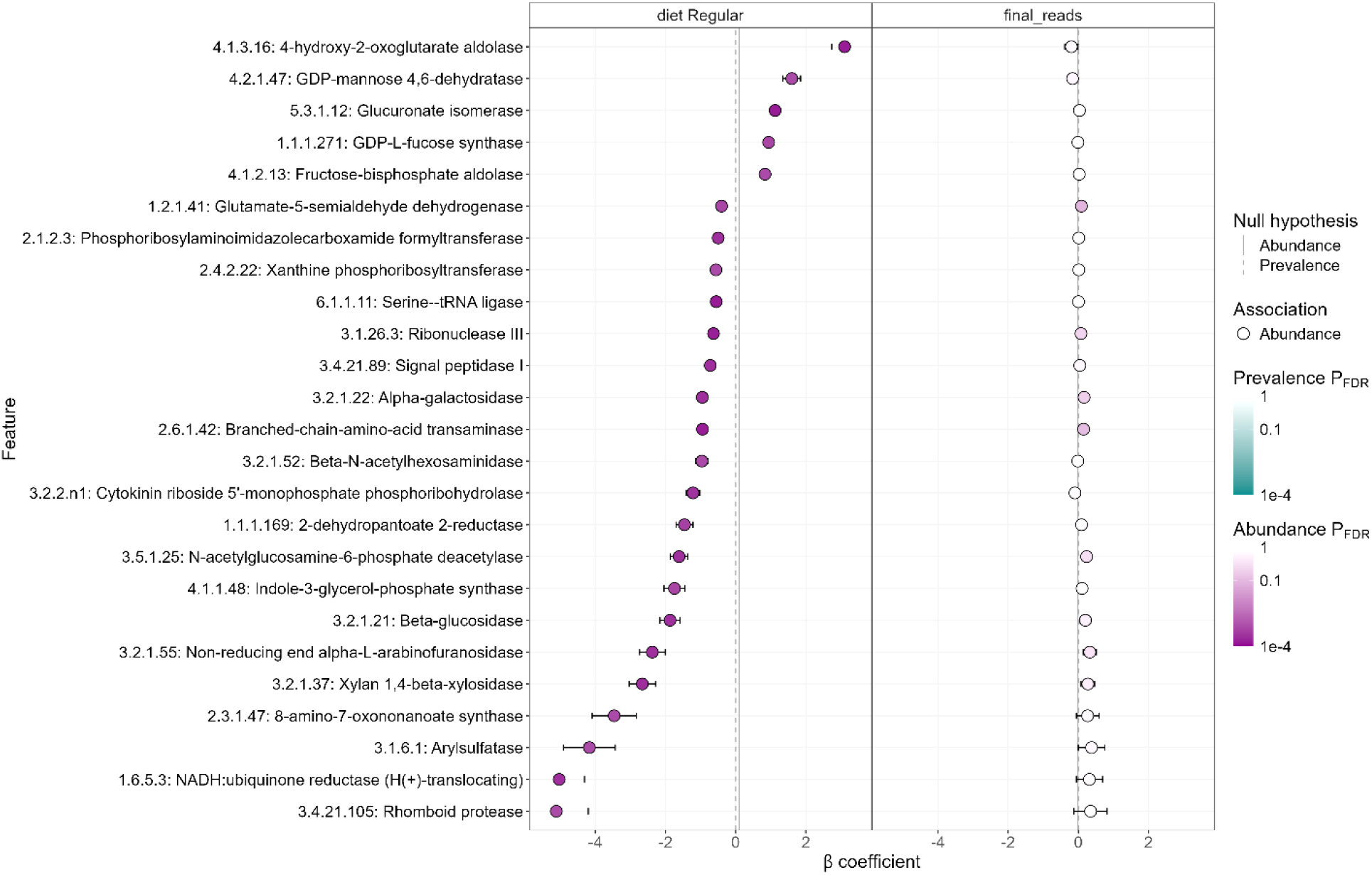
Microbial enzyme functions associated with diet, identified with MaAsLin 3. Coefficient plot for the 25 enzyme functions, labelled by Enzyme Commission (EC) number (profiled with [HUMAnN 3), with the strongest diet associations in a multivariable MaAsLin 3 model. The model included diet ([Regular vs. reference diet]) and sequencing depth (final_reads). Each row is one enzyme function, ordered by its diet coefficient.

**Supplementary Figure 5.**
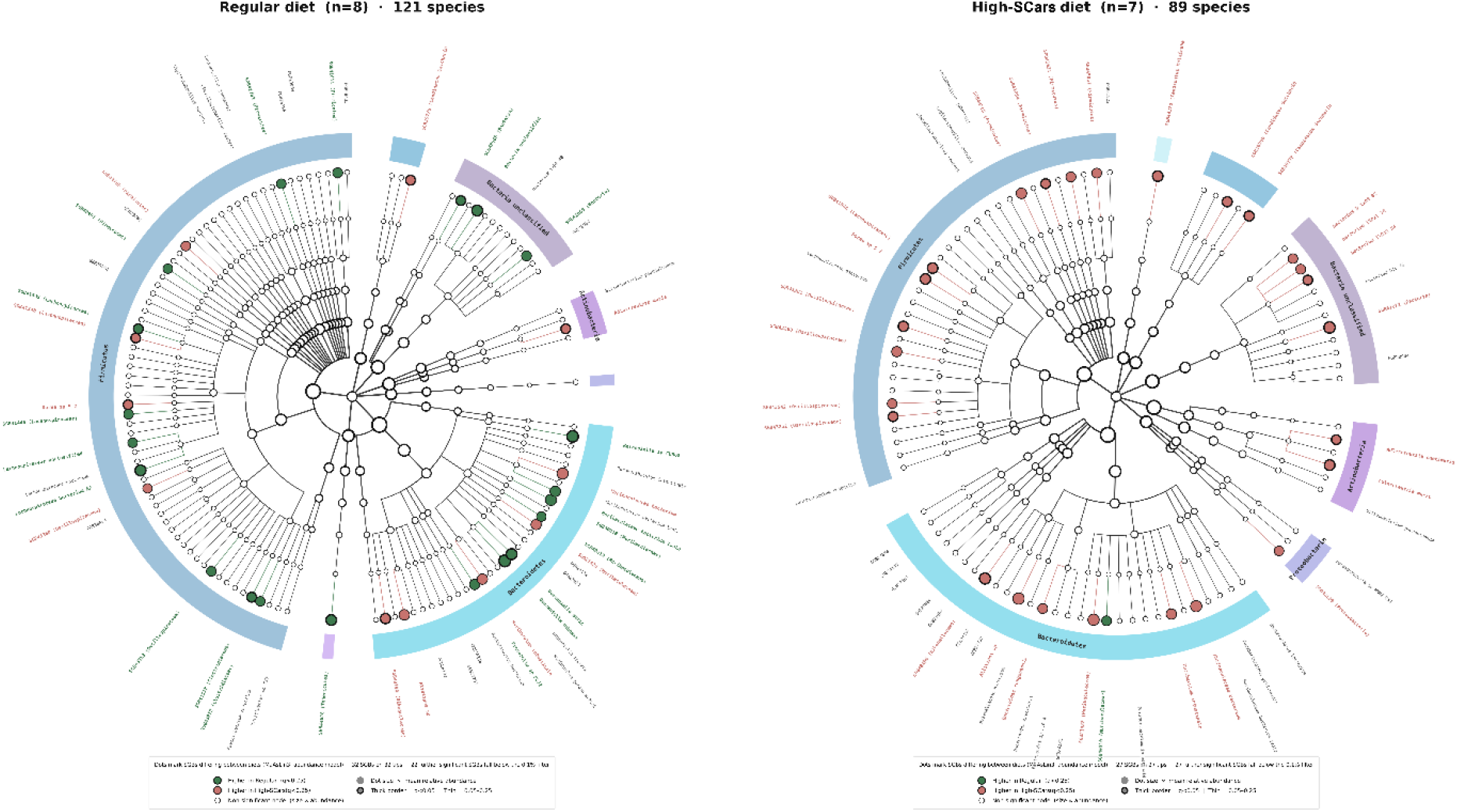
Inferred hierarchical taxonomic relationships (profiled with MetaPhlAn 4.0.6, ChocoPhlAn database: Oct22) between fecal microbiomes from Regular and High-SCars diet in circular cladogram designed using python codes generated by Claude AI (Sonnet 4.6).

